# Deciphering the evolution of the transcriptional and regulatory landscape in human placenta

**DOI:** 10.1101/2020.09.11.289686

**Authors:** Ming-an Sun, Gernot Wolf, Yejun Wang, Sherry Ralls, Anna D. Senft, Jinpu Jin, Caitlin E. Dunn-Fletcher, Louis J. Muglia, Todd S. Macfarlan

## Abstract

In mammals, the placenta mediates maternal-fetal nutrient and waste exchange and provides immunomodulatory actions that facilitate maternal-fetal tolerance. The placenta is highly diversified among mammalian species, yet the molecular mechanisms that distinguish the placenta of human from other mammals are not fully understood. Using an interspecies transcriptomic comparison of human, macaque, and mouse term placentae, we identified hundreds of genes with lineage-specific expression – including dozens that are placentally-enriched and potentially related to pregnancy. We further annotated the enhancers for different human tissues using epigenomic data and demonstrate that the placenta and chorion are unique in that their enhancers display the least conservation. We identified numerous lineage-specific human placental enhancers, and found they are highly overlapped with specific families of endogenous retroviruses (ERVs), including MER21A, MER4A/B and MER39B that were previously linked to immune response and placental function. Among these ERV families, we further demonstrate that MER41 insertions create dozens of lineage-specific Serum Response Factor (SRF) binding loci in human, including one adjacent to *FBN2*, a placenta-specific gene with increased expression in humans that produces the peptide hormone placensin to stimulate glucose secretion and trophoblast invasion. Our results demonstrate the prevalence of lineage-specific human placental enhancers which are frequently associated with ERV insertions and likely facilitated the lineage-specific evolution of the mammalian placenta.

## Introduction

The placenta is an extraembryonic organ essential for fetal development by facilitating maternal-fetal nutrient and waste exchange and providing immunomodulatory actions that facilitate maternal-fetal tolerance (Maltepe and Fisher 2015). As a transient tissue that is discarded after birth, the placenta has unique features including the tumor-like invasion of the maternal uterus (Beaman, et al. 2016) and the ability to escape from maternal immune attack (Ander, et al. 2019). Despite its common origin in all mammals, the placenta is extraordinarily diverse in its morphology, function and molecular features among mammalian species (Chuong, et al. 2010; Schmidt, et al. 2015; Roberts, et al. 2016). For example, the mature placenta of mouse has a yolk sac structure that degenerates after the first trimester in human placenta (Schmidt, et al. 2015; Ross and Boroviak 2020), and the chorioallantoic placenta found in mouse and human are structurally distinct (Rossant and Cross 2001). Remarkable differences between the placenta of human and other primates have also been documented, for example the human-specific B KIR haplotypes and Siglec-6 and IMUP-2 expression (Schmidt, et al. 2015). These morphological and molecular differences limit the insights that animal models can provide for human pregnancy. Clarifying the molecular features unique to the human placenta is of great importance for better understanding of human development and evolution.

The immense diversification of mammalian placentae is reflected molecularly at the level of species-specific proteins and regulatory elements. One striking example is the lineage-specific co-option of endogenous retroviral envelope genes as “syncytin” genes, which are essential for trophoblast cell fusion (Mi, et al. 2000) and probably related to maternofetal tolerance (Mangeney, et al. 2007). These exapted syncytin genes have been identified in lineages including humans, mice, cows, sheep and rabbits, but not in pigs and horses which do not form a multinucleated syncytiotrophoblast layer (Furukawa, et al. 2014; Denner 2016). Endogenous retroviruses (ERVs) are one class of Transposable elements (TEs) (that also include LINEs, SINEs, and DNA transposons) that make up more than half of human genome and have been documented as critical facilitators of genome evolution (Feschotte 2008; Chuong, et al. 2016a). Co-option of TEs, in particular ERVs, seem to have been especially important for placenta evolution, since several other placental-specific genes, such as *PEG10* and *RTL1*, are also co-opted from ERVs (Rawn and Cross 2008). In addition to producing functional proteins, ERVs can also be co-opted as alternative promoters and enhancers (Cohen, et al. 2009). For example, the placental-specific transcription of *CYP19A1*, which controls estrogen synthesis, is driven by an alternative promoter created by MER21A insertion during primate evolution (van de Lagemaat, et al. 2003).

Many TEs inherently possess transcription factor binding sites and can quickly accumulate across the genome via a copy-and-paste mechanism, allowing TEs to be adopted as lineage-specific regulatory elements during evolution (Feschotte 2008). Lineage-specific TE insertions have been reported to facilitate the evolution of different regulatory networks including those for innate immunity (Chuong, et al. 2016b), p53 binding (Wang, et al. 2007), embryonic development (Kunarso, et al. 2010; Macfarlan, et al. 2012; Fuentes, et al. 2018) and pregnancy (Lynch, et al. 2011; Chuong, et al. 2013). Previous comparisons of mouse and rat trophoblast stem cells (TSCs) suggest that species-specific TSC enhancers are highly enriched for ERVs, with RLTR13D elements contributing to hundreds of mouse-specific enhancers bound by core trophoblast factors Cdx2, Eomes and Elf5 (Chuong, et al. 2013). Our previous study suggests that an anthropoid primate-specific insertion of a THE1B family ERV element controls the placental expression of *CRH* and alters gestation length (Dunn-Fletcher, et al. 2018). TEs were also reported to transform the uterine regulatory landscape and transcriptome during the evolution of mammalian pregnancy (Lynch, et al. 2015). However, a comprehensive assessment of how the transcriptional and regulatory elements evolve in human placenta and how TEs contribute to this evolutionary process is still lacking.

In this study, we performed a systematic interspecies comparison of the placental transcriptomes and epigenetically-defined putative enhancers among human, macaque and mouse. We identified hundreds of lineage-specific placental genes and numerous lineage-specific placental enhancers, which may underly the species-specific characteristics of human placenta. By providing a comprehensive view of the transcriptional and regulatory landscape in human placenta relative to other mammals, we demonstrate the prevalence of lineage-specific human placental enhancers that are frequently associated with ERV insertions. These ERV-derived enhancers likely facilitated the rapid diversification of placenta that is present in all mammals and crucial for pregnancy success.

## Results

### Interspecies transcription alterations among human, macaque and mouse placentae

The human lineage separated from Rhesus Macaque (hereby abbreviated as macaque) about 25 Million Years Ago (MYA), and primates and mice diverged about 100 MYA (**Figure 1A**). To determine the interspecies divergence of gene expression in placenta, we compared human, macaque and mouse transcriptomes of term placenta and other tissues (brain, heart, kidney, liver, testis) generated in this study or retrieved from public resources (**Table S1**). A computational procedure was implemented for interspecies gene expression analysis (**Figure S1**). In brief, we first quantified gene expression levels as TPM (Transcript Per Million). Then, we determined the 1:1:1 orthologs among the three species, which resulted in 14,023 protein-coding orthologs. Subsequently, the expression levels for these orthologs were normalized across interspecies samples and used for differential expression analysis and visualization. In agreement with previous findings (Brawand, et al. 2011), the interspecies samples are grouped by tissue instead of species, and human and macaque usually group more closely within each tissue (**Figure 1B, S2**).

We next compared the placentae of human, macaque and mouse to detect differentially expressed genes (DEGs). We identified 378 significant DEGs between human and mouse placentae – including 194 genes with increased expression in human (**Figure 1C, S3A, Table S2**). Many of these DEGs are known to be important for placenta: *CRH* and *CYP19A1* in particular were reported to have evolved expression in primate placenta (van de Lagemaat, et al. 2003; Dunn-Fletcher, et al. 2018) – the former encodes corticotropin-releasing hormone which controls the timing and onset of labor (Wadhwa, et al. 1998; Dunn-Fletcher, et al. 2018), and the latter encodes aromatase P450 to catalyze estrogen synthesis (Kamat, et al. 2002; van de Lagemaat, et al. 2003). Gene Ontology (GO) enrichment analysis demonstrated that human-placenta-enriched genes are associated with extracellular region, receptor binding, growth factor activity and female pregnancy (**Table S3**). A comparison of macaque vs. mouse placenta identified 95 DEGs, with 58.9% (n=56) also differentially expressed between human vs. mouse (**Figure 1C, S3B, Table S2**). The placental gene expression alterations between human vs. mouse and macaque vs. mouse are highly correlated (**Figure 1D**; Pearson’s r = 0.706, p-value<2.2e-16). As expected, inspection of the expression patterns of human vs. mouse DEGs further demonstrated that macaque placenta gene expression more closely resembles human (**Figure 1E**).

**Figure 1.**
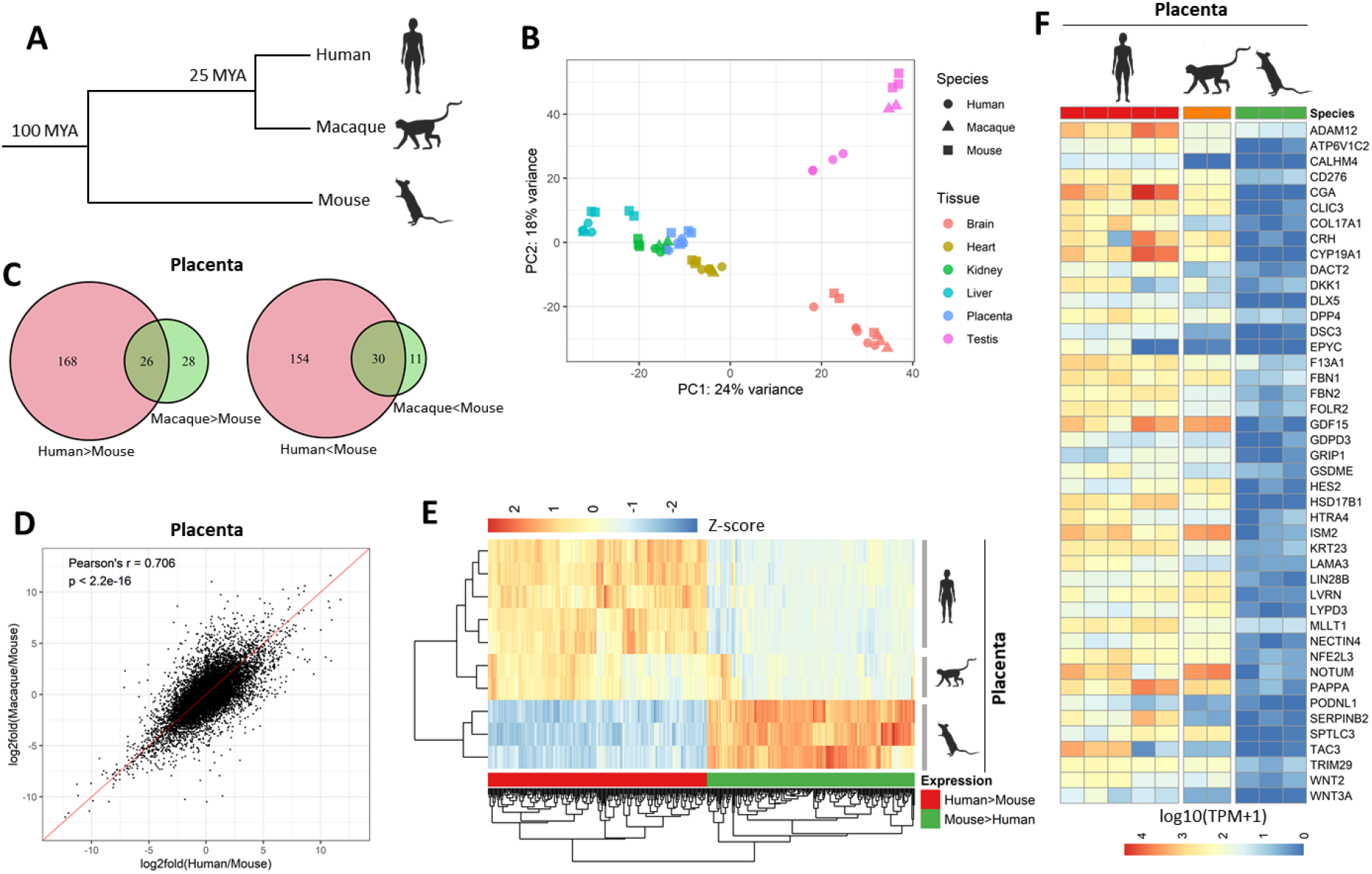
Global gene expression alterations among human, mouse and macaque placentae. **A**. Diagram shows the divergence time among human, macaque and mouse. **B**. Relationship of different human, macaque and mouse samples based on PCA results. Top 5000 genes with the highest expression variance were used for PCA. The species and tissues are indicated by shape and color, respectively. **C**. Overlap of DEGs identified between Human vs. Mouse and Macaque vs. Mouse. Human>Mouse indicates DEGs with increased expression in human, and Human<Mouse for DEGs with decreased expression in human. Macaque>Mouse and Macaque<Mouse can be interpreted similarly. **D**. Positive correlation of placental gene expression alterations in human and macaque relative to mouse. The x-axis and y-axis indicate the log2fold(Human/Mouse) and log2fold(Macaque/Mouse) of the normalized TPM values for the 1:1:1 orthologs. The red line indicates the linear fitting result. The Pearson’s r and p-value are indicated. **E**. Expression pattern of the DEGs identified between human and mouse placentae in the placentae of three species. The color gradient indicates z-score. **F**. Expression patterns of the 44 DEGs that show both increased expression in human against mouse as well as enriched expression in placenta compared to 16 other human tissues from BodyMap 2.0. The color gradient indicates log10(TPM+1) values.

We further asked how many DEGs between human and mouse placentae are specifically expressed in placenta. To answer this question, we compared the transcriptome of human placenta against 16 other human tissues from BodyMap 2.0 (**Figure S4**). While thousands of genes (range from 3,395 to 6,334) with placenta-enriched expression can be identified relative to other specific tissues, the number dropped sharply when comparing against multiple tissues simultaneously (**Figure S5**). As a result, we identified 311 genes with placenta-enriched expression relative to all 16 tissues (**Figure S6, Table S4**) – supporting a previous assessment that the number of truly placenta-specific genes is relatively small (Rawn and Cross 2008). These genes include many known placental genes (Rawn and Cross 2008), including *CRH, ENDOU, PAPPA, PEG10, PGF, PHLDA2, PRL, TAC3* and Pregnancy Specific Glycoprotein genes (**Table S4**). GO enrichment analysis demonstrates these placenta-specific genes are highly associated with extracellular region, female pregnancy, viral envelope and hormone activity (**Table S5**). Several enriched GO terms (eg. extracellular region and female pregnancy) are also associated with genes with increased expression in human placenta relative to mouse (**Table S3**). Furthermore, for DEGs with increased expression in human placenta relative to mouse, almost one-fourth (44/194) have placenta-enriched expression (**Figure 1F**).

### Interspecies transcriptional alterations correlate with changes of histone modifications in promoter regions

The covalent modifications of histone tails are widely used to annotate the activity of regulatory elements (Xiao, et al. 2012). For example, H3K4me3 is used for annotating active promoters, H3K27ac for active enhancers and promoters, and H3K4me1 for active/poised enhancers and promoters (Piunti and Shilatifard 2016). To unravel the epigenetic mechanisms underlying the evolution of gene expression among species, we examined the occupancy of these three histone modifications among human, macaque and mouse placentae using ChIP-Seq (**Table S1**). We first calculated their normalized intensity in promoter regions, which were defined as +/− 1 kb flanking transcript start sites (TSSs). Three groups of genes with Human>Mouse, Human<Mouse or unchanged expression in placenta were ranked based on log2foldChange (**Figure 2A**), and then the interspecies changes of each histone modification in the corresponding promoter regions were plotted alongside the ranked genes (**Figure 2B-D**). We demonstrate that the interspecies gene expression alterations between human vs. mouse placentae are accompanied by changes in histone modifications in the promoter regions of these DEGs. This correlation is most evident for H3K27ac and H3K4me3 which mark active regulatory elements (**Figure 2B-D, S7**). Further comparison of the macaque and mouse placentae shows that the histone modifications change similarly in macaque and human (**Figure 2E-G**), in agreement with the observation that gene expression patterns in human and macaque placentae are highly similar (**Figure 1C-F**).

**Figure 2.**
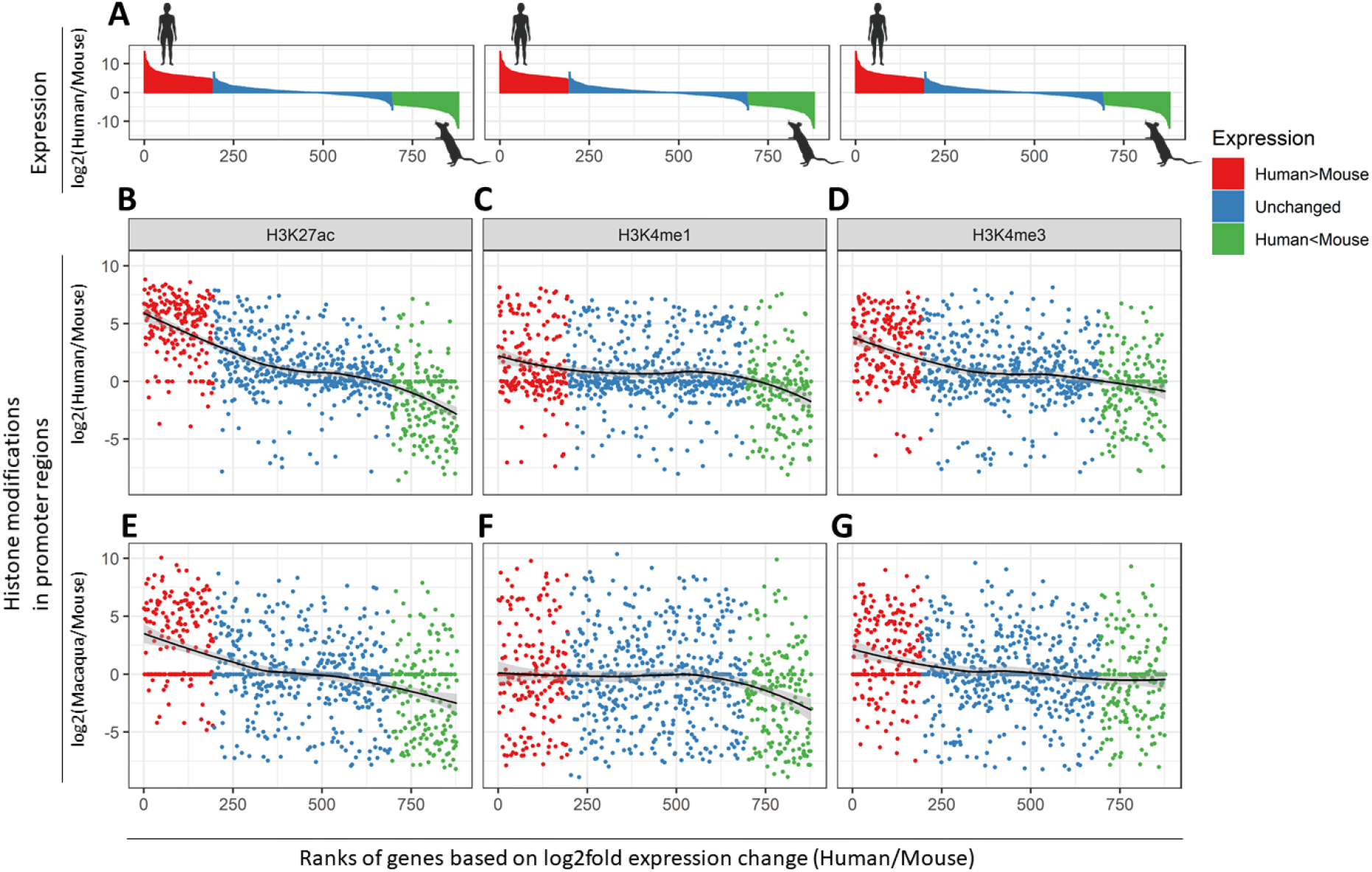
Interspecies transcriptional alterations are correlated with changes of histone modifications in promoter regions. **A**. Expression alterations for genes with Human>Mouse, Human<Mouse and unchanged expression between human and mouse placentae. For genes with unchanged expression, only 500 randomly selected genes are shown. The genes within each group are ranked by log2fold expression change between human and mouse placentae. The x-axis indicates the gene rank, and y-axis indicates log2fold expression change based on normalized TPM values. **B-D**. Interspecies alterations of H3K27ac (**B**), H3K4me1 (**C**) and H3K4me3 (**D**) in promoter regions (+/− 1 KB of TSSs) between human and mouse placentae. Similar to **A**, the x-axis indicates the rank of genes based on log2fold expression changes. The y-axis indicates log2fold changes of histone modification in promoter regions between human and mouse placentae. Smoothed curves were calculated to indicate the trend. **E-G**. Similar to **B-D**, except that the y-axis is for the histone modification changes between macaque and mouse placentae.

### Identification of numerous lineage-specific enhancers in human placentae

Enhancers play a central role for the precise spatiotemporal regulation of gene expression (Long, et al. 2016). Since placenta is amongst the most diversified organs in mammals (Maltepe and Fisher 2015), we reasoned placental enhancers may be even less conserved than those for other tissues. To test this, we retrieved the H3K4me3 and H3K27ac peaks for human placenta and 14 other human tissues from the ENCODE project (**Table S1**). In all 15 tissue together we annotated 301,370 putative enhancers as H3K27ac(+) H3K4me3(–) regions located more than 500 bp away from TSSs, and further annotated the enhancers specific for each tissue. As expected, enhancers for placenta and chorion (a crucial part of the fetal part of the placenta) show strikingly low conservation relative to other tissues, for example the spinal cord, where enhancers are highly conserved (**Figure 3A**, **S8**). We also used the LiftOver to match the human placental enhancers to the genomes of mouse and macaque and found that a large fraction of them are absent from the mouse genome. This is most striking for placenta and chorion, where about half of human enhancers cannot be matched to mouse genome (**Figure S9A**). Further, even though most human enhancers can be matched to the macaque genome, placenta and chorion still have the highest percentage of lineage-specific enhancers relative to other tissues (**Figure S9B**) – together suggesting that placental/chorion enhancers are more rapidly evolving.

**Figure 3.**
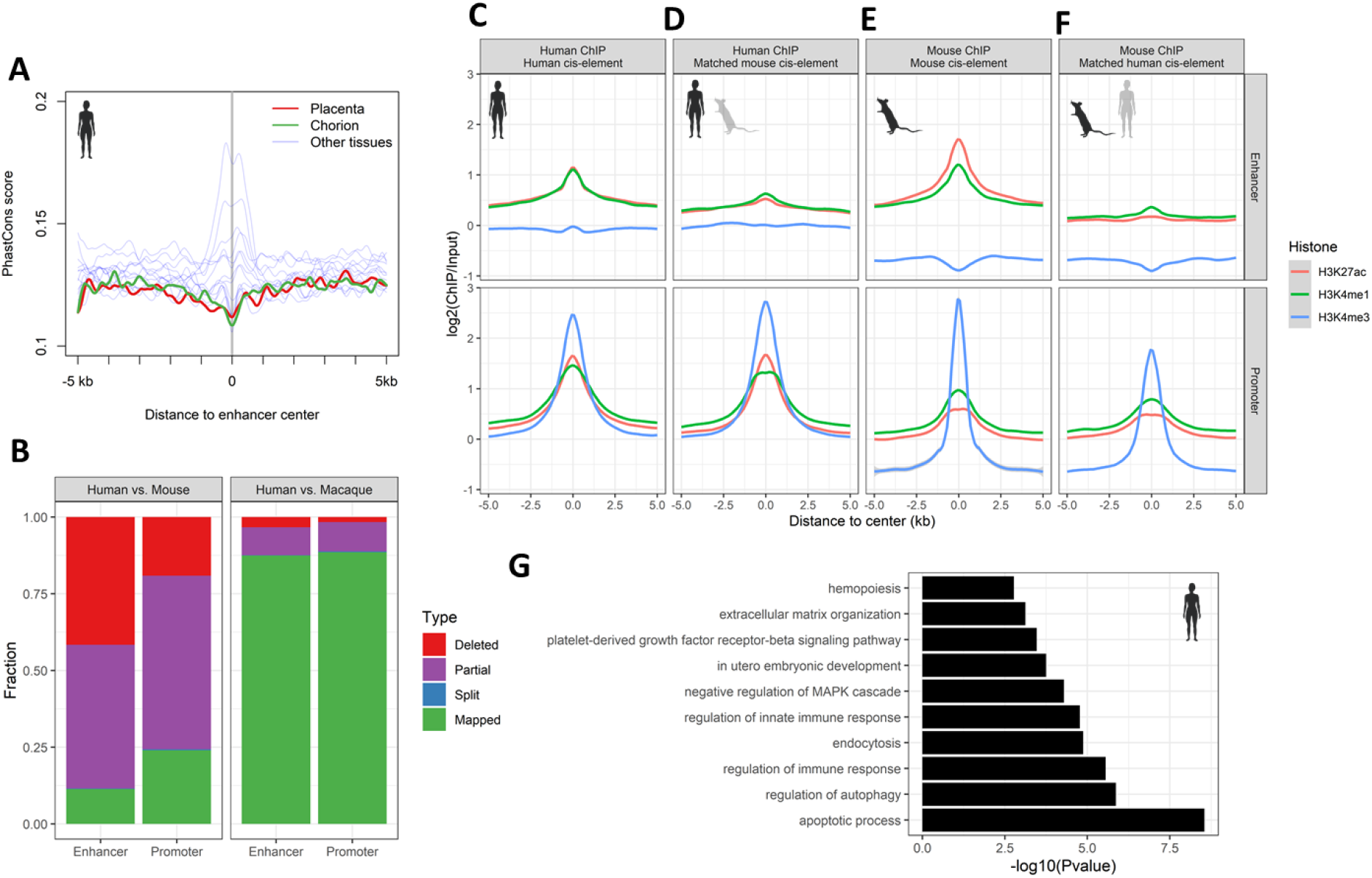
Interspecies evolution of enhancers among human, macaque and mouse placentae. **A**. Sequence conservation flanking the enhancers specific for placenta, chorion and 13 other human tissues (refer to **Figure S8** and **Figure S9** for more details). The y-axis indicates averaged phastCons score. The curves for placenta and chorion are highlighted in red and green respectively, while other tissues (adrenal, aorta, kidney, liver, lung, ovary, pancreas, spinal cord, spleen, stomach, thymus, thyroid, uterus) are in grey. Cubic spline was used for curve smoothing. **B**. Mapping of human placenta enhancers and promoters to mouse and macaque genomes, respectively. Mapped, split, partial and deleted regions are based on LiftOver matching results and indicated with different colors. The y-axis shows the fraction for each group of enhancers or promoters. **C-F**. The intensity of H3K27ac, H3K4me1 and H3K4me3 flanking enhancers and promoters annotated based on the same data (**C,E**) or matching from other species by LiftOver (**D,F**). **G**. Enriched GO terms for lineage-specific human placental enhancers. Only 10 representative GO terms from “Biological process” category are shown.

To further examine the conservation of human placental enhancers and promoters, we annotated each using the newly generated H3K27ac and H3K4me3 data. We annotated 86,968 enhancers and 28,581 promoters in human placenta. By matching against the mouse genome using LiftOver, we demonstrated that 41.6% (n = 36,217) of human placental enhancers are absent from the mouse genome, which is much higher than the 19.1% for promoters (**Figure 3B**) – suggesting that a large fraction of human placental enhancers are newly acquired during evolution. This raises the question: for human placental regulatory elements that can be matched to the mouse genome, are they also active in mouse placenta? We examined mouse genomic intervals matching human placental enhancers and found that they usually lack the active histone marks H3K4me3 and H3K27ac – in contrast to mouse genomic intervals matching human placental promoters which have comparable levels of active histone marks (**Figure 3C-F**). Lineage-specific human placental enhancers are associated with apoptotic processes, immune response, endocytosis and in utero embryonic development (**Figure 3G**). Taken together, we demonstrated placenta enhancers are rapidly evolving among the compared species both at sequence and activity levels.

### Specific ERV families contribute to interspecies evolution of human placental enhancers

TEs, and in particular LTR retrotransposons (primarily ERVs), have been reported to facilitate the evolution of immune system and placenta (Chuong, et al. 2013; Chuong, et al. 2016b; Dunn-Fletcher, et al. 2018). After identifying thousands of lineage-specific human placental enhancers, we further asked if specific groups of TEs contribute to the evolution of these enhancers. We first examined the occurrence of major classes of TEs, and found that LTR, SINE, LINE and DNA transposons are significantly overrepresented in lineage-specific human placental enhancers (**Figure 4A**). Among them, LTR retrotransposons are enriched to the highest degree (Enrichment fold = 4.4, p-value < 2.2e-16), which were thus examined further. Using a Binomial Test-based approach, we identified 42 LTR/ERV families (Bonferroni-adjusted p-value<0.01) that are significantly overrepresented in lineage-specific human placenta enhancers (**Figure 4B-C, Table S6**). Interestingly, several of them were previously reported to be associated with placenta development or the immune system. For example, MER41A/B was shown to facilitate innate immunity evolution (Chuong, et al. 2016b), and MER21A and MER39B were previously linked to placenta (Emera, et al. 2012; Pavlicev, et al. 2015). Inspection of the H3K27ac occupancy surrounding the most significantly enriched families showed that H3K27ac occupancy usually centers on the overlapped ERV elements (**Figure 4D**).

**Figure 4.**
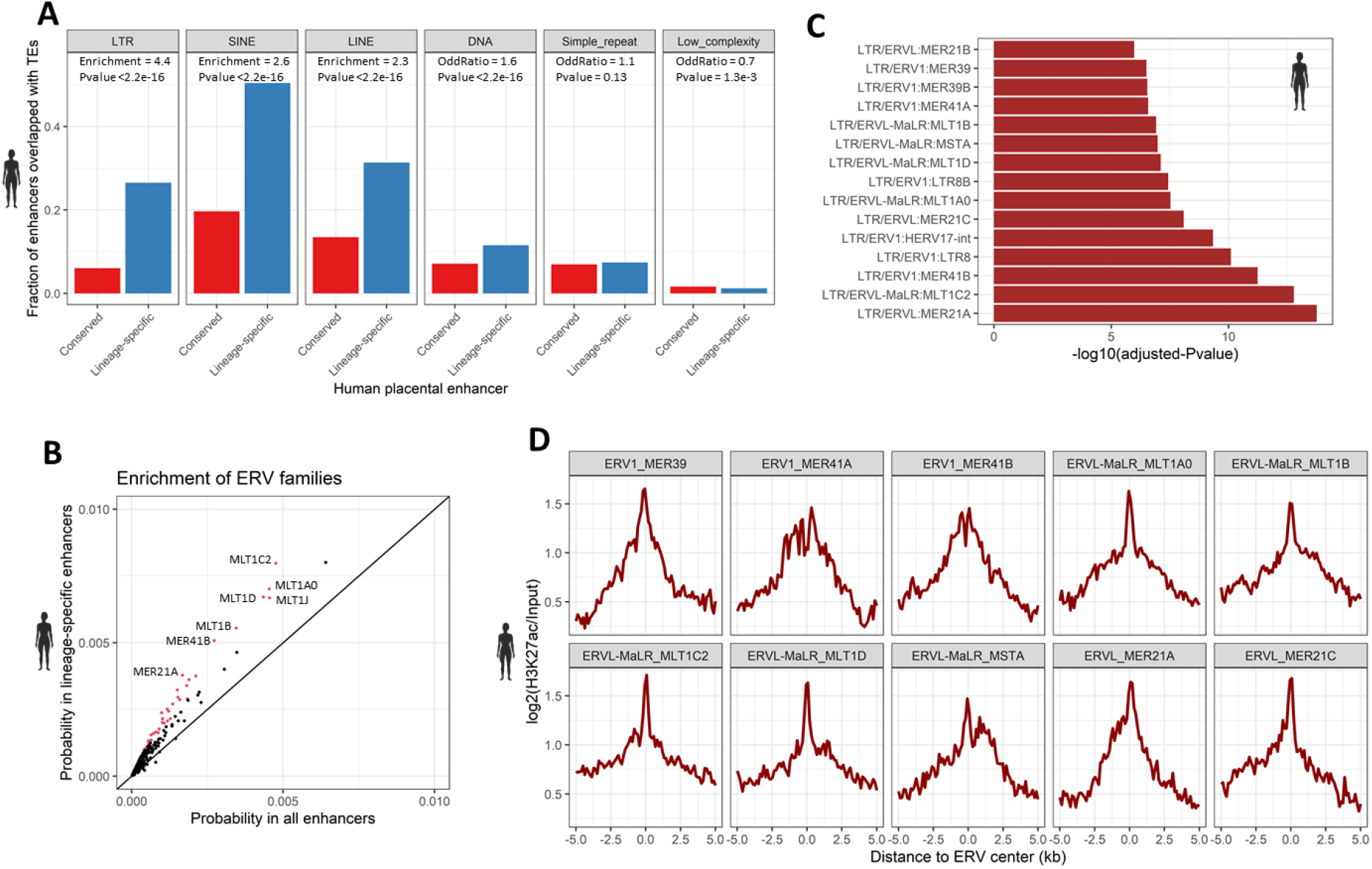
Contribution of ERV families for the interspecies evolution of human placental enhancers. **A**. Enrichment of major classes of TEs in lineage-specific human placental enhancers. The y-axis shows the fraction of enhancers overlapped with TEs. The conserved (ie. mapped) and lineage-specific (ie. deleted) enhancers are annotated based on LiftOver matching to mouse genome. The enrichment of each class of TEs in lineage-specific enhancers relative to conserved enhancers are tested by Fisher’s Exact Test, with the enrichment fold and p-values indicated. **B**. Enrichment of ERV families in lineage-specific human placental enhancers. The x-axis and y-axis indicate the fraction of all enhancers and lineage-specific enhancers overlapped with each ERV family. Each dot represents one ERV family, with significantly enriched ERV families (Bonferroni-adjusted p-value<0.01) highlighted in red. **C**. Representative ERV families that are highly overrepresented in lineage-specific human placental enhancers. The x-axis indicates −log10(adjusted-pvalue). Only the top 15 ERV families are shown. **D**. H3K27ac intensity surrounding the center of ERV elements that overlapping with lineage-specific human placental enhancers. ERV families with at least 100 overlapped elements are shown. The curves are calculated based on ERVs located within lineage-specific human placental enhancers.

### MER41 insertion events create lineage-specific SRF binding loci in human

Many LTR retrotransposons harbor transcription factor binding motifs, thus their insertions in the genome provide a shortcut for creating lineage-specific enhancers. One example is MER41, which facilitates human innate immunity evolution by creating primate-specific STAT1 binding motifs (Chuong, et al. 2016b). MER41 elements are also overrepresented within lineage-specific human placental enhancers (**Figure 4C**). Motif enrichment analysis demonstrated that in addition to STAT1, binding motifs for several other transcription factors (eg. SP8, ZN350, ANDR and SRF) are also highly enriched in MER41B elements within lineage-specific human placental enhancers (**Figure 5A, Table S7**). Among them, *SRF* (Serum response factor) is a transcription factor that regulates actin genes for cytoskeleton maintenance and immediate early genes for cellular response (Miano, et al. 2007). In response to a variety of extracellular signals in the serum, SRF binds the serum response elements near its target genes to stimulate their expression (Clark and Graves 2014). Even though the role of SRF in placenta remains unclear, increasing evidence suggests SRF is involved in immune response (Taylor, et al. 2014), cell proliferation, invasion and adhesion (Franco, et al. 2013; Gualdrini, et al. 2016) and angiogenesis (Franco, et al. 2008; Franco, et al. 2013) – processes also important for placenta function. Further, forced expression of SRF in mouse TSCs was shown to promote the differentiation into giant cells (Asanoma, et al. 2007). Therefore, we further examined whether SRF binds in MER41 elements.

**Figure 5.**
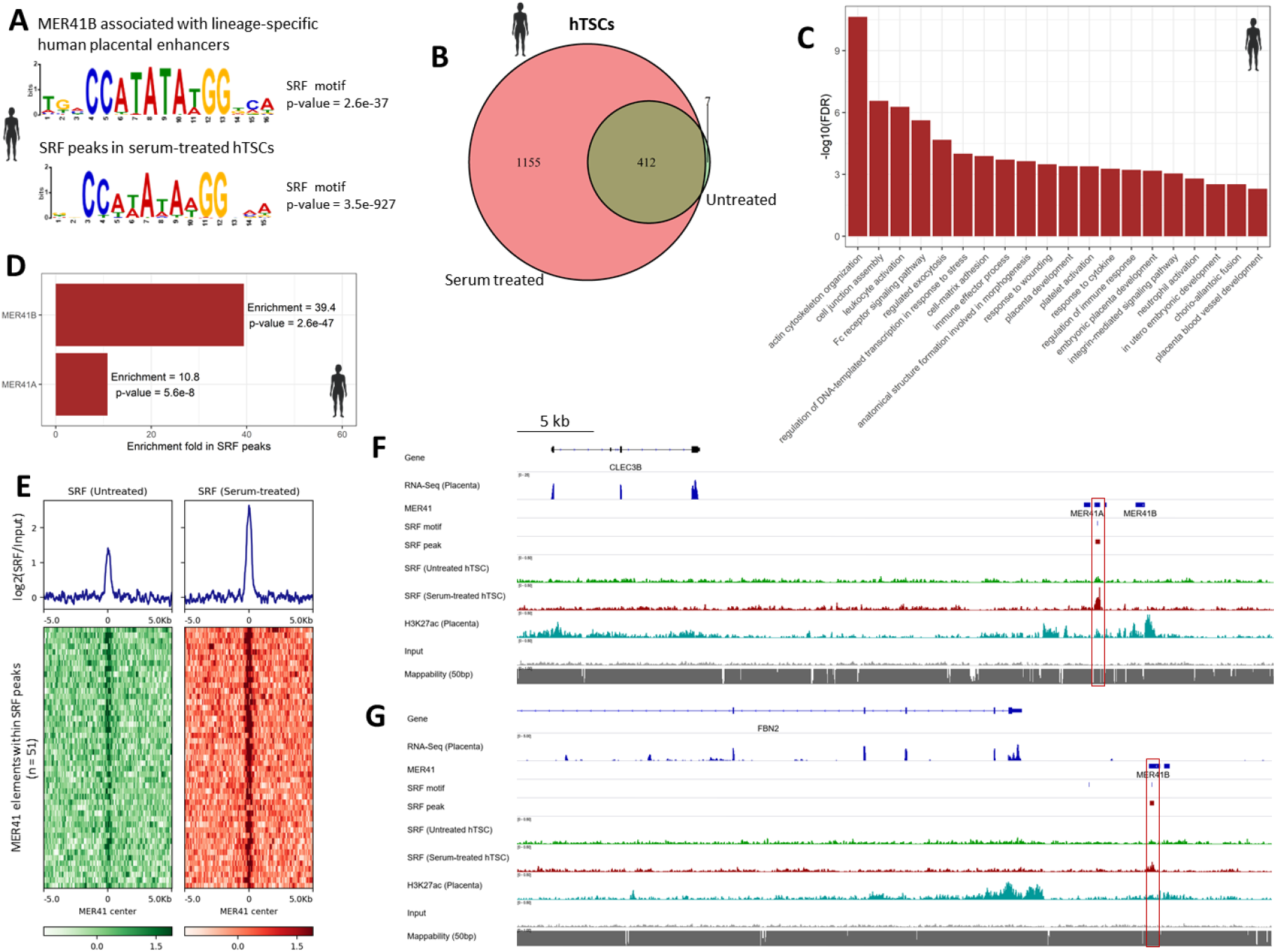
MER41 insertions create lineage-specific SRF binding loci in human. **A**. SRF motifs enriched in MER41B associated lineage-specific human placental enhancers (top) and SRF peaks in serum-treated hTSCs (bottom). **B**. Venn diagram showing the overlap of SRF peaks identified from untreated and serum-treated hTSCs. **C**. GO terms associated with SRF peaks identified from serum-treated hTSCs. Only 20 representative GO terms from the “Biological Process” category are shown. **D**. Enrichment of MER41A and MER41B elements in SRF peaks from serum-treated hTSCs. The enrichment fold and p-values are labelled. **E**. SRF binding flanking the MER41 elements that occur within SRF peaks in hTSCs. The averaged curves (top) and heatmaps (bottom) are shown – all are centered on the center of the 51 MER41 elements within SRF peaks identified from serum-treated hTSCs. **F,G**. IGV tracks showing the expression, SRF motif occurrence, SRF binding and H3K27ac occupancy of the two MER41-SRF peaks associated genes CLEC3B (**F**) and FBN2 (**G**). The MER41-associated SRF peaks are highlighted in red square.

Human trophoblast stem cells (hTSCs) have been successfully established recently (Okae, et al. 2018) and serve as a promising *in vitro* model for studying human placenta development. We applied ChIP-Seq to profile the genome-wide binding of SRF before and after serum treatment in hTSCs. Inspection of several canonical SRF target genes (*ACTB, JUNB, EGR1* and *EGR2*) confirmed the data quality (**Figure S10**). We identified 420 SRF peaks in untreated hTSCs, and 1,567 after serum treatment (**Figure 5B**). The canonical SRF motif – which resembles the one overrepresented in MER41B elements within lineage-specific human placental enhancers – is significantly enriched in SRF binding loci in serum-treated hTSCs (**Figure 5A**). Apart from its expected function in actin cytoskeleton organization and cell-matrix adhesion, we found that SRF binding loci are also significantly associated with immune regulation, placenta development and chorio-allantoic fusion (**Figure 5C, Table S8**). Interestingly, SRF binding loci are associated with phenotypes including embryo size, prenatal body size, placenta vasculature and labyrinth morphology, and T cell/B cell physiology (**Table S9**) – together suggest that SRF is potentially involved in placental development and immunity in human.

We further examined SRF binding in different groups of MER41 elements. Our results demonstrate that after serum treatment, SRF binding on MER41B is significantly increased (**Figure S11**). MER41A elements are also bound by SRF yet with lower intensity, while other MER41 groups (MER41C, MER41D, MER41E, MER41G and MER41-int) do not show any sign of SRF binding. Consistently, MER41A and MER41B elements are overrepresented within SRF peaks, with enrichment fold of 10.8 and 39.4 respectively (**Figure 5D**). We further demonstrate that for MER41-associated SRF peaks, SRF binding is centered on the MER41 elements (**Figure 5E**) – suggesting that SRF recognizes MER41 elements. Interestingly, for genes with TSSs less than 50 kb away from MER41-SRF peaks, the majority have increased expression in human placenta relative to mouse (**Figure S12**). In particular, *CLEC3B* and *FBN2* display significant interspecies expression alterations between human and mouse. Manual inspection showed that MER41-associated SRF peaks near these two genes in serum-treated hTSCs have evident H3K27ac occupancy in human placenta (**Figure 5F,G**). Both *FBN2* and *CLEC3B* have placenta-enriched expression (**Figure S13**), and interestingly *FBN2* was recently reported to secrete a peptide hormone, placensin, that promotes placental cell invasiveness (Yu, et al. 2020). Together, our results suggest that specific MER41 elements may facilitate the expression of some human placenta specific genes by creating SRF-bound enhancers.

## Discussion

The placenta is an organ with extraordinary diversity in mammals (Chuong, et al. 2010). To clarify the molecular mechanisms underlying human placenta evolution, we performed interspecies transcriptomic and epigenomic comparison to examine the alterations of the transcriptional and regulatory landscape among the placentae of human, macaque and mouse. We demonstrate that placenta and chorion have the least conserved enhancers compared with other human tissues. We further demonstrate that specific families of ERVs are significantly associated with lineage-specific human placental enhancers. These ERVs include several families (eg. MER21A, MER41A/B, MER39B) that were previously linked to placental function or immune response. These results indicate that the prevalence of lineage-specific placental enhancers, which are frequently created by lineage-specific ERV insertions, may underlie the extreme diversification of mammalian placentae.

MER41 is one family of ERVs known to regulate human innate immunity by creating lineage-specific STAT1 binding sites (Schmid and Bucher 2010; Chuong, et al. 2016b). We found that MER41 elements associated with lineage-specific human placental enhancers are also enriched for the binding motif of SRF, which is originally known for regulating actin genes and immediate early genes (Miano, et al. 2007) and more recently found to also control immune response (Taylor, et al. 2014), cell proliferation, invasion and adhesion (Franco, et al. 2013; Gualdrini, et al. 2016) and angiogenesis (Franco, et al. 2008; Franco, et al. 2013). Since these processes are important for placental function, we examined SRF binding in human TSCs. Indeed, SRF-bound regions are associated with placenta-related functions, such as placenta development, in utero embryonic development, chorio-allantoic fusion and placenta blood vessel development. We demonstrate that SRF binds dozens of MER41A/B elements, and genes adjacent to MER41-associated SRF peaks usually have increased expression in the placenta of human relative to mouse. Specifically, the associated gene *FBN2* is a placenta-specific gene recently reported to secrete the peptide hormone placensin which promotes placental cell invasiveness (Yu, et al. 2020). Notably, *FBN2* is highly expressed in the placenta of human but not mouse (Yu, et al. 2020), and higher degree of trophoblast invasion is a remarkable feature for human placenta relative to other species (Cohen and Bischof 2007). These results indicate the possible contribution of MER41 elements in driving the evolution of invasive potential of human placenta by creating lineage-specific enhancers.

While we systematically examined the interspecies molecular alterations in human placenta on both the transcriptional and regulatory level, our study may suffer from several limitations. First, the interspecies expression comparison is focused on 1:1 orthologs, which preclude the possibility to examine the gained, lost or duplicated genes for human placenta evolution. Second, while we demonstrate that MER41-associated SRF peaks occur near placental genes with human-increased expression such as *FBN2* and *CLEC3B*, currently it is difficult to further validate the function of these putative enhancers due to lack of mature methods for manipulation of hTSCs. Third, since the transcriptomic and epigenomic data are not profiled at single-cell resolution, it is uncertain how much of the detected interspecies transcriptional and regulatory alterations are due to differences in cell population. Finally, most enhancers are distal from promoters, we therefore could not establish the relationship of many putative human placental enhancers to their target genes. Despite of these limitations, this study provides the most comprehensive picture to date about the transcriptional and regulatory landscape evolution in human placenta, which is valuable for our understanding about human placenta evolution. Importantly, the identified genes and regulatory elements with evolved expression/activity also provide valuable candidates for further studies examining human placenta evolution.

## Materials and methods

### Placenta sample collection

Placenta samples were collected and processed as described in a previous study (Dunn-Fletcher, et al. 2018). Briefly, two human placentae were collected by cesarean section at 39 weeks 1 day and 39 weeks 2 days gestational age (IRB protocol: CCHMC IRB 2013–2243). Two macaque placentae were collected by cesarean section at 128 days and 131 days gestational age (80% completed gestation). Collected placenta samples were frozen in −80°*C* for later use.

### Human trophoblast stem cell culture

hTSCs derived from human cytotrophoblast cells (Okae, et al. 2018) were a gift from the Okae lab. They were cultured following a previous protocol as described (Okae, et al. 2018). In brief, hTSCs were grown in trophoblast stem cell medium (TSM) consisting of DMEM/F12 (Gibco, 1263401) supplemented with 2 mM Glutamax (Fisher, 35050061), 0.1 mM b-Mercaptoethanol (Gibco, 21985-023), 0.2% FBS (HyClone, SH3007103), 0.5% Pen-Strep (Fisher, 15140122), 0.3% BSA (Sigma-Aldrich, A9576-50ml), 1% ITS-X (Fisher, 51500056), 1.5 μg/mL L-ascorbic acid (Sigma-Aldrich, A92902-100G), 50 ng/mL hEGF (Fisher, 10605HNAE25), 2 μM CHIR99021 (Sigma-Aldrich, SML1046-5MG), 0.5μM A83-01 (Sigma-Aldrich, SML0788-5MG), 1 μM SB431542 (Sigma-Aldrich, 616464-5MG), 0.8 mM VPA (Wako, 227-01071) and 5 μM Y27632 (Abcam, ab120129). Several reagents, including hEGF, CHIR99021, A83-01, SB431542, VPA, Y27632 and L-ascorbic acid, were added fresh before use. hTSCs were cultured on Collagen-IV (Corning, CB-40233) coated TC dishes at 37 °C with 5% CO_2_ and split at 1:2 to 1:4 ratio every two days. Serum treatment was performed by culturing hTSCs in TSM mixed with 15% FBS (HyClone, SH3007103) for 30 min.

### RNA-Seq

Total RNA for placenta and hTSC samples was extracted using RNeasy Micro kit (Qiagen, 74004) with on-column DNase digestion. RNA samples were submitted for library construction by TruSeq stranded mRNA sample preparation kit (Illumina) and sequencing as 75 bp paired-end reads with HiSeq2500 (Illumina). We also collected RNA-Seq data for dozens of human, macaque and mouse tissues from public resources (**Table S1**). Reads were trimmed with Trim Galore v0.6.4 (https://github.com/FelixKrueger/TrimGalore). To perform differential expression analysis of the samples from the same species, we aligned trimmed reads to the corresponding reference genome using STAR v2.7.1 (Dobin, et al. 2013), and then obtained gene-level read counts using the featureCount function from subread v2.0.0 (Liao, et al. 2013). Finally, differentially expressed genes were identified using DESeq2 v1.28.0 (Love, et al. 2014) with the cutoff: FDR<0.05 and foldchange>2. Notably, when both paired-end and single-end data are available for compared samples, only data for read 1 are used.

To perform differential expression analysis among different species, we first calculated Transcripts per Million (TPM) value for each gene using RSEM v1.3.2 (Li and Dewey 2011) with reference genome and gene annotation downloaded from ENSEMBL database (Yates, et al. 2020). Taking the calculated TPM values as input, interspecies differential expression analysis was performed using ExprX (https://github.com/mingansun/ExprX). In brief, it first determined the 1: 1 orthologs between compared species based on the homolog annotations from ENSEMBL database (Yates, et al. 2020). The 1:1 orthologs were then filtered to only keep protein-coding genes after excluding ribosomal genes and genes from chromosomes X, Y and MT. Finally, the expression of the filtered 1:1 orthologs were normalized using TMM method (Robinson, et al. 2010), and differential expression analyses between compared species were performed using Rank Product method (Hong, et al. 2006) with FDR<0.05 and log2foldchange>1 as cut-off.

### ChIP-Seq

ChIP-Seq for placenta samples was performed using the MAGnify^TM^ Chromatin Immunoprecipitation System (ThermoFisher, 492024), and ChIP-Seq for hTSC samples was performed following previous protocol (Blecher-Gonen, et al. 2013) with minor modifications. Chromatin fragmentation was performed using Diagenode Bioruptor Plus Sonicator. The antibodies used include H3K4me1 (Abcam, ab8895), H3K4me3 (Abcam, ab8580), H3K27ac (Abcam, ab4729) and SRF (ActiveMotif, 61385). The amount of chromatin used for each reaction is 30 μg for transcription factors and 20 μg for histone modification. ChIP-Seq libraries were constructed using Takara SMARTer ThruPLEX DNA-Seq Kit (Takara, R400674), and then sequenced as 50 bp single-end reads with HiSeq2500 (Illumina).

Reads were trimmed with TrimGalore v0.6.4 and then aligned to the corresponding reference genome using Bowtie v2.3.5 (Langmead and Salzberg 2012) with default settings. PCR duplicates were removed using the rmdup function of samtools v1.10 (Li, et al. 2009). The data reproducibility between biological replicates was confirmed, then reads from replicates were pooled together for further analysis. Peak calling was performed with MACS v2.2.5 (Zhang, et al. 2008) with default settings. H3K4me1 was called as broad peaks, while H3K4me3, H3K27ac and SRF were called as narrow peaks. The peaks for human and mouse were further cleaned by removing those overlapped with ENCODE Blacklist V2 regions (Amemiya, et al. 2019).

### Reference genome and annotation

Reference genome and gene annotation for human (GRCh38), macaque (Mmul_10) and mouse (GRCm38) were downloaded from ENSEMBL database (release 100) (Yates, et al. 2020). PhastCons annotation for human, macaque and mouse were downloaded from UCSC Genome Browser (Lee, et al. 2020). Transposable element annotations for human were downloaded from the RepeatMasker website (http://www.repeatmasker.org/) on May 27, 2016. The clades for ERV families were obtained from Dfam database (Hubley, et al. 2016).

### Gene Ontology and motif enrichment analysis

Gene Ontology enrichment analyses for differentially expressed genes were performed using DAVID functional annotation tools (Huang da, et al. 2009). Gene Ontology enrichment analyses for genomic regions (eg. ChIP-Seq peaks and putative enhancers) were performed with GREAT (McLean, et al. 2010). Motif enrichment analysis was performed using MEME-ChIP (Machanick and Bailey 2011), with sequences +/− 500 bp from the center of ChIP-Seq peaks or ERV elements as input.

### Regulatory element annotation and comparison

To identify human-placenta-enriched enhancers, we first inferred the enhancers for human, macaque and mouse placenta respectively based on histone modifications. Putative Enhancers were defined as H3K27ac peaks that lack H3K4me3 occupancy and not within +/− 500 bp from TSSs. The initially called putative enhancers are usually of very different length, which may influence the mapping among species. Thus, each enhancer was adjusted as +/− 100 bp from its center. Finally, length-adjusted placental enhancers were compared among species using UCSC LiftOver (Lee, et al. 2020) with default settings.

### Association of ERV families to lineage-specific human placental enhancers

To determine if certain ERV families are overrepresented within lineage-specific human placental enhancers, we designed a Binomial-Test based approach. In brief, we first calculated the occurrence of each group of ERV family within all enhancers and within lineage-specific enhancers using the windowBed function of BEDtools (Quinlan and Hall 2010). The occurrence of each family in all enhancers were counted and considered as the background. Then, Binomial Test was performed to determine if an ERV subfamily is significantly overrepresented within lineage-specific human placental enhancers relative to all human placental enhancers. To control for Family-Wise Error Rate, the calculated p-values were further adjusted (Bonferroni correction).

### Statistical analysis and data visualization

All statistical analyses were performed with R statistical programming language (Team 2016). The enrichment of MER41 elements within SRF peaks was tested using the fisher function from BEDtools (Quinlan and Hall 2010). Heatmaps for ChIP-Seq data were generated using DeepTools (Ramirez, et al. 2014). Heatmaps from gene expression clustering analysis were generated using pheatmap (https://github.com/raivokolde/pheatmap). PCA analysis and visualization were performed using the plotPCA function in DESeq2 (Love, et al. 2014). RNA-Seq and ChIP-Seq tracks were visualized using IGV (Thorvaldsdottir, et al. 2013).

### Data deposit

All the data generated in this study were deposited in NCBI Gene Expression Ominibus (GEO) under accession number GSE153082, GSE153083 and dbGap under accession number phs002233.

## Acknowledgements

This work was supported by the Intramural Research Program of the NICHD, NIH and the Priority Academic Program Development of Jiangsu Higher Education Institutions (PAPD). LJM received support through the March of Dimes Prematurity Research Center Ohio Collaborative and the NIH/NICHD (HD 091527). We thank Dr. Suhas Kallapur for assistance with the rhesus samples. We thank the National Institute of Child Health and Human Development (NICHD) Molecular Genomics Core for high-throughput sequencing. This study utilized the computational resources of the NIH High-Performance Computing Biowulf cluster (https://hpc.nih.gov) and Yangzhou University College of Veterinary Medicine High-Performance Computing cluster.

